# Soluble amyloid-beta buffering by plaques in Alzheimer disease dementia versus high-pathology controls

**DOI:** 10.1101/227009

**Authors:** Thomas J. Esparza, Mihika Gangolli, Nigel J. Cairns, David L. Brody

**Affiliations:** Department of Neurology, Washington University, St. Louis, Missouri, United States of America; Department of Biomedical Engineering, Washington University, St. Louis, Missouri, United States of America; Knight Alzheimer Disease Research Center, Washington University, St. Louis, Missouri, United States of America; Hope Center for Neurological Disorders, Washington University, St. Louis, Missouri, United States of America

## Abstract

An unanswered question regarding Alzheimer disease dementia (ADD) is whether amyloid-beta (Aβ) plaques sequester toxic soluble Aβ species early in the pathological progression. We previously reported that the concentration of soluble Aβ aggregates from patients with mild dementia was higher than soluble Aβ aggregates from patients with modest Aβ plaque burden but no dementia. The ratio of soluble Aβ aggregate concentration to Aβ plaque area fully distinguished these groups of patients. We hypothesized that initially plaques may serve as a reservoir or sink for toxic soluble Aβ aggregates, sequestering them from other targets in the extracellular space and thereby preventing their toxicity. To initially test a generalized version of this hypothesis, we have performed binding assessments using biotinylated synthetic Aβ_1-42_ peptide. Aβ_1-42_-biotin peptide was incubated on unfixed frozen sections from non-demented high plaque pathology controls and patients with dementia of the Alzheimer type. The bound peptide was measured using ELISA and confocal microscopy. We observed no quantitative difference in Aβ binding between the groups using either method. Further testing of the buffering hypothesis using various forms of synthetic and human derived soluble Aβ aggregates will be required to definitively address the role of plaque buffering as it relates to ADD.

## Introduction

The relationship between Aβ plaque pathology and Alzheimer disease dementia (ADD)(1) has been a topic of considerable controversy. Soluble Aβ aggregates have been more directly linked to toxicity *in vitro* and in some animal model systems (2). We recently reported that soluble Aβ aggregates (termed “oligomers” in prior publications) were elevated in aqueous brain lysates from patients with early ADD in comparison with lysates from patients with Aβ plaque pathology but no dementia (3). However, despite statistically significant differences, there was still considerable overlap between groups. In further analyses, we found that the *ratio* of soluble Aβ aggregate levels to plaque area instead fully distinguished between patients with early ADD from patients with plaques but no dementia. This finding was replicated in a second cohort of cases. While it is likely that this finding indicates a fundamentally important pathophysiological linkage underlying ADD, the interpretation of the result is not straightforward. We posited in our initial report that plaques could serve as binding reservoirs or buffers for soluble Aβ aggregates; at early stages, Aβ plaques could adequately buffer soluble aggregates, protecting the nearby neuropil from toxicity, whereas at later times if buffering capacity was lost or overwhelmed, soluble aggregates could be free to diffuse in the extracellular space and exert toxicity (3). Direct binding and unbinding of soluble aggregates to the plaques themselves would be the most straightforward mechanism, but binding and unbinding to peri-plaque elements including dystrophic neurites, microglia, and astrocytes would be functionally equivalent. This idea was also based in part on the observations of Koffie et al., who reported loss of synapses in a gradient around plaques both in transgenic mice (4) and human brain sections (5), along with a halo of immunoreactivity consistent with soluble aggregates/oligomeric species; these results were interpreted as consistent with release of synaptotoxic soluble aggregates from plaques. Because synapse loss correlates strongly with dementia (6, 7) the failure of buffering leading to synaptic toxicity could be proposed as a substrate of dementia. The question of whether failure of buffering of soluble Aβ by plaques underlies ADD leads to several non-mutual exclusive hypotheses (2):

> Hypothesis 1: <u>Qualitative</u> changes in the Aβ plaques correlate with dementia severity; plaques from non-demented controls retain more buffering capacity than plaques from demented patients.
>
> Hypothesis 2: <u>Qualitative</u> changes in the <u>soluble Aβ aggregates</u> correlate with dementia; soluble Aβ aggregates from demented cases bind to Aβ plaques less avidly than soluble Aβ aggregates from non-demented controls.
>
> Hypothesis 3: <u>Quantitative</u> changes in the <u>soluble Aβ aggregates</u> correlate with dementia; intrinsic buffering properties of soluble Aβ aggregates and plaques from non-demented controls are similar to those of demented patients when assessed using the same concentrations of soluble Aβ aggregates, only the quantity of the soluble aggregates changes with progression of the disease.

Here we report our first efforts towards testing Hypothesis 1. Specifically, we assessed Aβ buffering capacity of plaques in frozen brain slices from non-demented controls vs. patients with early dementia. While we have made initial progress in purifying soluble Aβ aggregates from human brain tissue (8), we have not yet achieved sufficient purification for to use native human brain aggregates for these buffering studies. Therefore, we used biotinylated synthetic Aβ_1-42_ peptide for these initial experiments, reasoning that if plaque buffering of soluble Aβ is a general property, that this approach could yield insight into the buffering properties.

## Materials and Methods

### Regulatory Compliance

All protocols were carried out in accordance with the Charles F. and Joanne Knight Alzheimer Disease Research Center, Washington University guidelines. This specific study was approved by the Knight Alzheimer Disease Research Center. All donors or family members gave informed consent for a brain donation and use in research studies.

### Selection and sectioning of human frontal cortical tissue

Clinically and neuropathologicaly well-characterized cases were obtained from the Knight Alzheimer Disease Research Center, Washington University School of Medicine, Saint Louis, Missouri, USA. Brain tissue was obtained from the frontal lobe and included the cortical ribbon and underlying white matter (Brodmann areas 8/9). Cognitive status was determined during life using the Clinical Dementia Rating (CDR). At the time of expiration a final CDR was ascertained using established procedures. Alzheimer pathology was assessed using the NIA-AA neuropathologic diagnostic criteria. The samples included: CDR1 cases (n = 10, 85.2±10.8 years at death, 16.1±6.3 hours post mortem interval, 8 females & 2 males) and CDR0 with Aβ pathology cases (n=6, 90.4±9.6 years at death, 11.7±8.5 hours post mortem interval, 4 female & 2 male).

Unfixed frontal cortex was embedded in ‘optimum cutting temperature’ (O.C.T., Sakura Finetek, USA) compound and frozen by immersion in liquid nitrogen chilled 2-methylbutane. The tissue blocks were then equilibrated at -20°C and 18μm sections were cut and mounted directly onto positively charged glass slides using a Leica CM 1950 cryostat. The tissue sections were stored at -80°C prior to use.

For quantification of Aβ plaque pathology, adjacent tissue samples were trimmed and placed into 4% paraformaldehyde for 48 hours and then transferred into 30% sucrose (w/v) in 1x phosphate buffered saline for an additional 48 hours. Free-floating 50μm sections were cut using a HM 430 sliding microtome (Thermo Fisher Scientific, Waltham, MA) and stored in a cryoprotectant solution (0.44M sucrose, 2.7M ethylene glycol, 30mM sodium phosphate buffer, pH 7.4) prior to immunohistochemistry.

### Immunohistochemical staining of human cortical tissue

Free-floating 50μm tissues sections were washed 3x in Tris-buffered saline (TBS) for 5 minutes each and then incubated with 0.3% H_2_O_2_ in TBS for 10 minutes at room temperature to block endogenous peroxidase. Following the incubation, sections were rinsed in TBS 3x for 5 minutes each, and then blocked with 5% normal goat serum (NGS) in TBS containing 0.25% (v/v) Triton X100 for 30 minutes at room temperature. Sections were then incubated with the Aβ-specific N-terminal mouse monoclonal HJ3.4 in 1% NGS in TBS-X at a 1:1000 dilution overnight at 4°C. The following day, sections were washed 3x in TBS for 5 minutes each and incubated with a biotinylated secondary goat anti-mouse antibody at a 1:1000 dilution in TBS-X for 1 hour at room temperature (Vector Laboratories). Following the incubation of the secondary antibody, the sections were washed 3x in TBS for 5 minutes each, incubated with ABC Elite (Vector Laboratories) at a 1:400 dilution in TBS for 1 hour at room temperature, then washed with TBS 3x for 5 minutes and developed with 3,3’-diaminobenzidine (#D5905; Sigma-Aldrich). Sections were mounted and dehydrated using a standard ethanol-xylene series followed by coverslipping. For each patient, three tissue sections separated by 600μm were stained for analysis.

### Quantification of immunohistochemical staining with ImageJ

Immunohistochemical samples from each patient were scanned using a Hamamatsu NanoZoomer HR model (Hamamatsu, Bridgewater, NJ). The percentage of gray matter containing HJ3.4 immunopositive Aβ plaque was measured using the Analyze Particles ImageJ (NIH) plug-in on thresholded 8-bit images with user defined gray-matter distinct regions of interest. The image thresholding was performed equally across the full sample set. During quantitation, the samples were coded such that the user was masked to the patient ID and CDR status.

### Binding of Aβ_1-42_-biotin to unfixed frozen human cortical tissue

For the binding experiments, unfixed frozen tissue sections (n=6 per patient) were equilibrated to ambient temperature. The residual O.C.T. compound was gently removed from surrounding the tissue and a hydrophobic barrier surrounding the tissue was prepared using an ImmEdge PAP pen (H-4000, Vector Laboratories, Burlingame, CA). Non-specific blocking of the tissue was performed using 200μl of a 1% bovine serum albumin (A7030, Sigma-Aldrich, St. Louis, MO) in artificial cerebral spinal fluid (aCSF) solution (148mM sodium chloride, 3mM potassium chloride, 1.4mM calcium chloride, 0.8mM magnesium chloride, 0.8mM sodium phosphate dibasic, 0.2mM sodium phosphate monobasic) for 1 hour at room temperature in a humidified chamber. The blocking solution was replaced with 150μl binding buffer (0.05% BSA in aCSF) containing 10nM N-terminal biotinylated synthetic Aβ_1-42_ (AS-23523, Anaspec, Fremont, CA) and incubated for 18 hours at room temperature in a humidified chamber. Following overnight binding, the tissue sections were gently washed 3x for 5 minutes each with binding buffer at room temperature. For direct quantification of the bound biointylated Aβ_1-42_ the tissue was dissociated using concentrated formic acid (n=3 sections/patient) and aliquoted into sterile microcentrifuge tubes. The formic acid in each sample aliquot was removed by vacuum centrifugation and the samples stored at -80°C until assayed. For confocal microscopy, the remaining sections (n=3/patient) were fixed with 4% paraformaldehyde in chilled methanol.

### Assessment of *ex-vivo* binding of Aβ_1-42_-biotin by confocal microscopy

The Aβ_1-42_-biotin bound then paraformaldehyde-fixed cortical tissue sections (n=3/patient) were subjected to signal amplification to allow for confocal image acquisition. The fixed sections were washed 3x in aCSF for 5 minutes each and incubated with ABC Elite at a 1:400 dilution in aCSF for 1 hour at room temperature. The signal was then amplified using the Biotin-XX Tyramide SuperBoost Kit (B40931, Thermo Fisher Scientific) with a development time of 10 minutes followed by washing 3x in aCSF for 5 minutes each. To quench endogenous autofluorescence the tissue sections were treated with TrueBlack (23007, Biotium, Fremont, CA) for 2 minutes followed by extensive washing in aCSF to remove residual reagent. The quenched, amplified sections were then incubated with streptavidin Alexa Fluor™ 594 conjugate (S11227, Thermo Fisher Scientific) at 1:1000 diluted in 0.1% BSA in aCSF for 30 minutes. The sections were washed 3x in aCSF for 5 minutes each, followed by counterstain with 4’,6-diamidino-2-phenylindole (DAPI, Sigma-Aldrich), and a final wash series before mounting with ProLong Gold Antifade media (P10144, Thermo Fisher Scientific). Images were acquired using an Olympus FV1200 scanning confocal microscope equipped with gallium arsenide phosphide detectors. Tiled (4x5), 7.5 micron z-stack images were acquired using a UPLFLN 20x (NA:0.70) objective with a laser output (0.5%) that produced non-saturated pixel data. All patient sections were acquired using the same instrument settings.

### Quantification of confocal imaging with ImageJ

Confocal acquired tiled images files (Olympus .OIB format) from each patient were imported into ImageJ for analysis. Using the DAPI channel, a global region of interest was defined to exclude non-tissue area. The Alexa Fluor™ 594 channel was converted to a z-projection using the “sum of slices” setting. A duplication of the z-projection was converted to an 8-bit image and used to create a threshold overlay mask within the Analyze Particles plugin. The overlay masks were imported into the ROI manager. The original z-projection was converted to an 8-bit image and the global ROI was used to deselect any overlay mask outliers. The global ROI and the overlay masks were used to measure the integrated density and area measures for each image. An additional measurement was made in the non-Alexa Fluor™ 594 region to serve as a background subtraction control. The image thresholding was performed equally across the full sample set. During quantitation, the samples were coded such that the user was masked to the patient ID and CDR status.

### Measurement of Aβ_1-42_ biotin by ELISA

Measurement of the formic acid soluble recovered N-terminally biotinylated Aβ_1-42_ was determined by ELISA using the mid-domain binding antibody HJ5.1 to capture and poly-streptavidin HRP-20 to detect bound peptide. HJ5.1 was coated to 96-well Nunc Maxisorp plates at 20 μ g/mL in carbonate buffer (35 mM sodium bicarbonate, 16 mM sodium carbonate, 3 mM sodium azide, pH 9.6) in 100 μl/well overnight at 4°C. Plates were washed 5x between steps with PBS using a BioTek EXL405 plate washer. The assay plates were blocked using 0.2 μm filtered 4% BSA in PBS for 1 hour at room temperature. Samples and standard were diluted in standard diluent, as described above, to a 100μL volume and loaded. An 8-point standard curve was generated using 5000, 1666.7, 555.6, 185.2, 61.7, 20.6, and 6.9 pg/mL of the N-terminal biotinylated synthetic Aβ_1-42_ loaded in triplicate. All samples were kept on ice during handling and the assay plates were incubated at 4°C overnight prior to development.

### Measurement of total Aβ_1-x_ by ELISA

The quantification of the Aβ_1-x_ isoforms was performed as previously described (8). The samples were resuspended in 5M guanidine-HCl prior to dilution with standard buffer (0.2 μm filtered 0.25% BSA, 0.5 M guanidine-HCl, 0.005% Tween-20, 300 mM Tris, 3 mM sodium azide, 2 μg/mL aprotinin, 1 μ g/mL leupeptin, in PBS). The samples were loaded onto HJ5.1 coated 96-well Nunc Maxisorp plates, which were blocked with 4% BSA, in addition to synthetic Aβ_1-40_ monomer standard loaded in triplicate. Following overnight incubation, the assay plates were detected using biotinylated HJ3.4 which binds N-terminally intact Aβ but should not bind to N-terminally biotinylated Aβ_1-42_. The assay was developed as previously described using poly-streptavidin HRP-20 (65R-S103PHRP, Fitzgerald, Acton, MA) and addition of 3,3’,5,5’-tetramethylbenzidine (T5569, Sigma-Aldrich) with the absorbance read on a BioTek Synergy 2 plate reader at 650 nm.

### Total protein quantification

Total protein was quantified in the formic acid dissociated samples using a fluorescence-based 96-well NanoOrange assay (N6666, Thermo Fisher Scientific). Sample aliquots were resuspended in 4M urea as well as dilutions of the reference BSA standard to normalize for effect on the assay. Appropriate dilutions for each samples and the standard were combined with assay reagent in a 96-well black microplate and measured by excitation at 485nm and emission at 590nm using a BioTek Synergy 2 plate reader.

### Assessment of Aβ_1-42_-biotin during binding by size-exclusion chromatography

To determine if qualitative size changes or quantitative loss of Aβ_1-42_-biotin occurred during the room-temperature overnight incubation period, we assessed pre- and post-incubation samples by size-exclusion chromatography. Clean slides without tissue were prepared and blocked the same as for the tissue sections. The blocking solution was replaced with 150μl binding buffer (0.05% BSA in aCSF) containing 10nM N-terminal biotinylated synthetic Aβ_1-42_ and incubated for 18 hours at room temperature in a humidified chamber. For the pre-incubation sample, the peptide solution was immediately recovered into a blocked-microcentrifuge tube. The recovered sample was injected onto a pre-equilibrated Superdex 200 10/300 GL column pre-equilibrated with 0.05% BSA in aCSF. The sample was eluted at 0.5 ml/min at 4°C and then immediately diluted and loaded onto an ELISA plate coated with HJ5.1 for quantitative analysis. The overnight incubation sample was recovered and assessed by size-exclusion chromatography in the same manner.

## Results

### Plaque coverage and morphology was indistinguishable between nondemented controls vs. patients with early ADD

As demonstrated in our previous report(3), Aβ plaques immunoreactive with the Aβ N-terminal specific antibody HJ3.4 covered similar portions of the gray matter in non-demented controls compared to patients with early ADD(**Fig. 1**). The brain samples used in these experiments came from different cases than those used for our previous report, but the demographics of the patients and the characteristics of the Aβ plaque pathologies were very similar (**Table 1**).

**Table 1:** Characteristics of human brain frontal cortex samples.

| Patient No. | Status | Age (yr.) | PMI (hr.) | Gender |
| --- | --- | --- | --- | --- |
| 1 | CDR 0 + path | 97 | 3.5 | M |
| 2 | CDR 0 + path | 91.7 | 12 | F |
| 3 | CDR 0 + path | 100.9 | 21 | F |
| 4 | CDR 0 + path | 95.4 | 23 | F |
| 5 | CDR 0 + path | 80.8 | 5.5 | M |
| 6 | CDR 0 + path | 76.7 | 5 | F |
| 7 | CDR 1 | 86.4 | 6.7 | F |
| 8 | CDR 1 | 89 | 19 | F |
| 9 | CDR 1 | 92.7 | 23 | F |
| 10 | CDR 1 | 94.2 | 11.6 | F |
| 11 | CDR 1 | 104.4 | 19 | F |
| 12 | CDR 1 | 68.6 | 21 | M |
| 13 | CDR 1 | 86 | 18 | F |
| 14 | CDR 1 | 81 | 21 | F |
| 15 | CDR 1 | 72.7 | 4.5 | M |
| 16 | CDR 1 | 76.8 | 17 | F |

**Figure 1:**
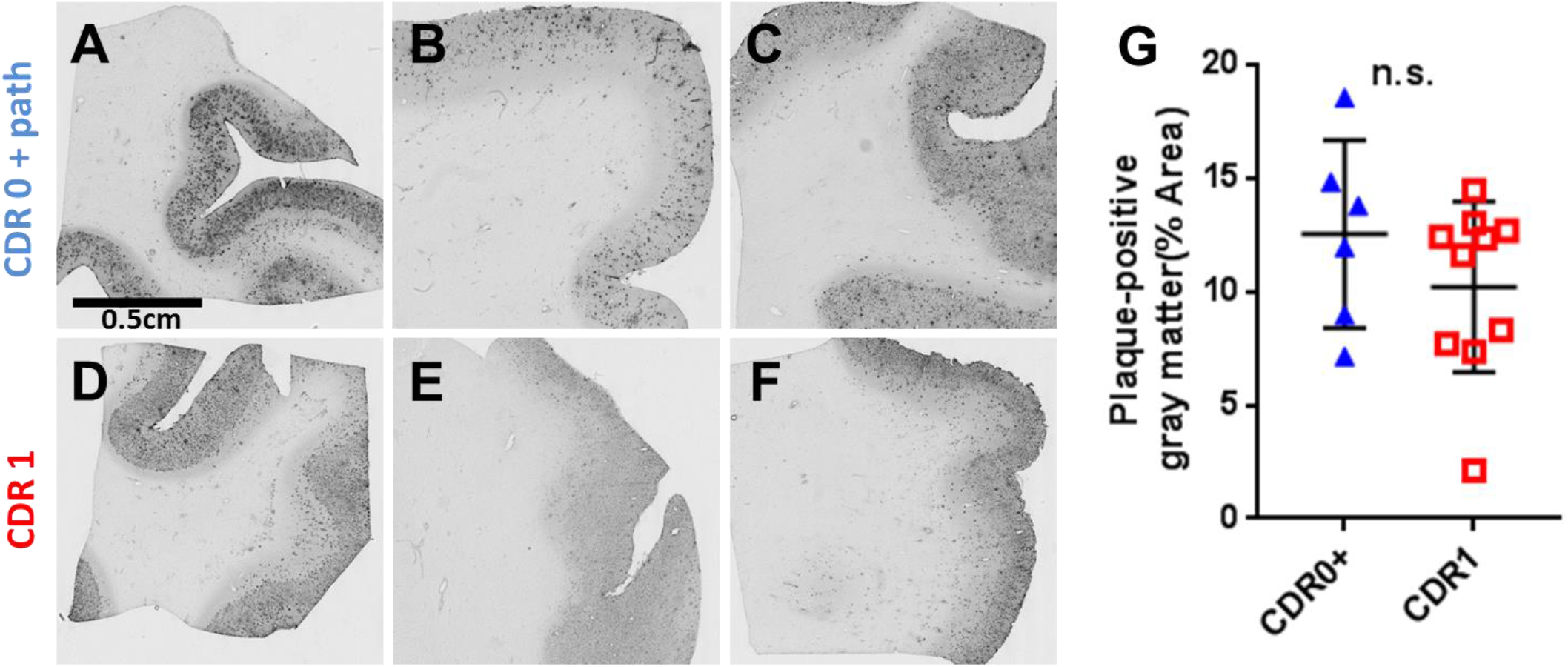
Representative amyloid-beta (Aβ) immunohistochemistry demonstrates similar levels of pathology between elderly high-pathology controls and elderly subjects with mild ADD. Scale bar = 0.5 cm, applies to panels A-F. (**A-C**) Aβ plaque pathology in frontal cortex sections from nondemented elderly subjects (CDR 0 + plaques). (**D-F**) Aβ plaque pathology in frontal cortex sections from elderly subjects with mild dementia of the Alzheimer type (CDR 1). (**G**) Gray matter coverage by Aβ plaque pathology was not different in the nondemented elderly subjects with plaques (CDR 0 + plaques) versus subjects with mild ADD (CDR 1) type (not significant [n.s.] by Mann–Whitney U test).

### No difference between binding of biotinylated Aβ_1-42_ bound to plaques in non-demented controls vs. plaques in patients with early ADD: results from confocal imaging

After binding of soluble biotinylated Aβ_1-42_ for 18 hours at room temperature to frozen sections followed by gentle washing in binding buffer (0.05% BSA in aCSF), biotinylated Aβ_1-42_ was retained in structures similar in size and morphology to Aβ plaques and did not appear to non-specifically bind throughout the tissue(**Fig 2A, B, D-F**). No binding was detected in brain sections from patients without plaques (**Fig 2C**), or on blank slides. Binding of biotinylated Af _42_ covered 0.407+/-0.124% of the brain area in slices from non-demented controls and 0.433+/ 0.089% of the area in slices from patients with early dementia. The area covered did not differ between groups (**Fig 3A**, p=0.6299). The area covered by bound soluble biotinylated Aβ_1-42_ di not correlate with the total immunohistochemically detected plaque coverage (**Fig 3B**). The total fluorescence intensity also did not differ between groups (p=0.9463). After normalization by total Aβ plaque area, total fluorescence intensity similarly did not differ between groups (**Fig 3C**). Thus, the binding capacity of plaques for synthetic biotinylated Aβ_1-42_ does not appear to differentiate demented from non-demented high plaque control brains.

**Figure 2:**
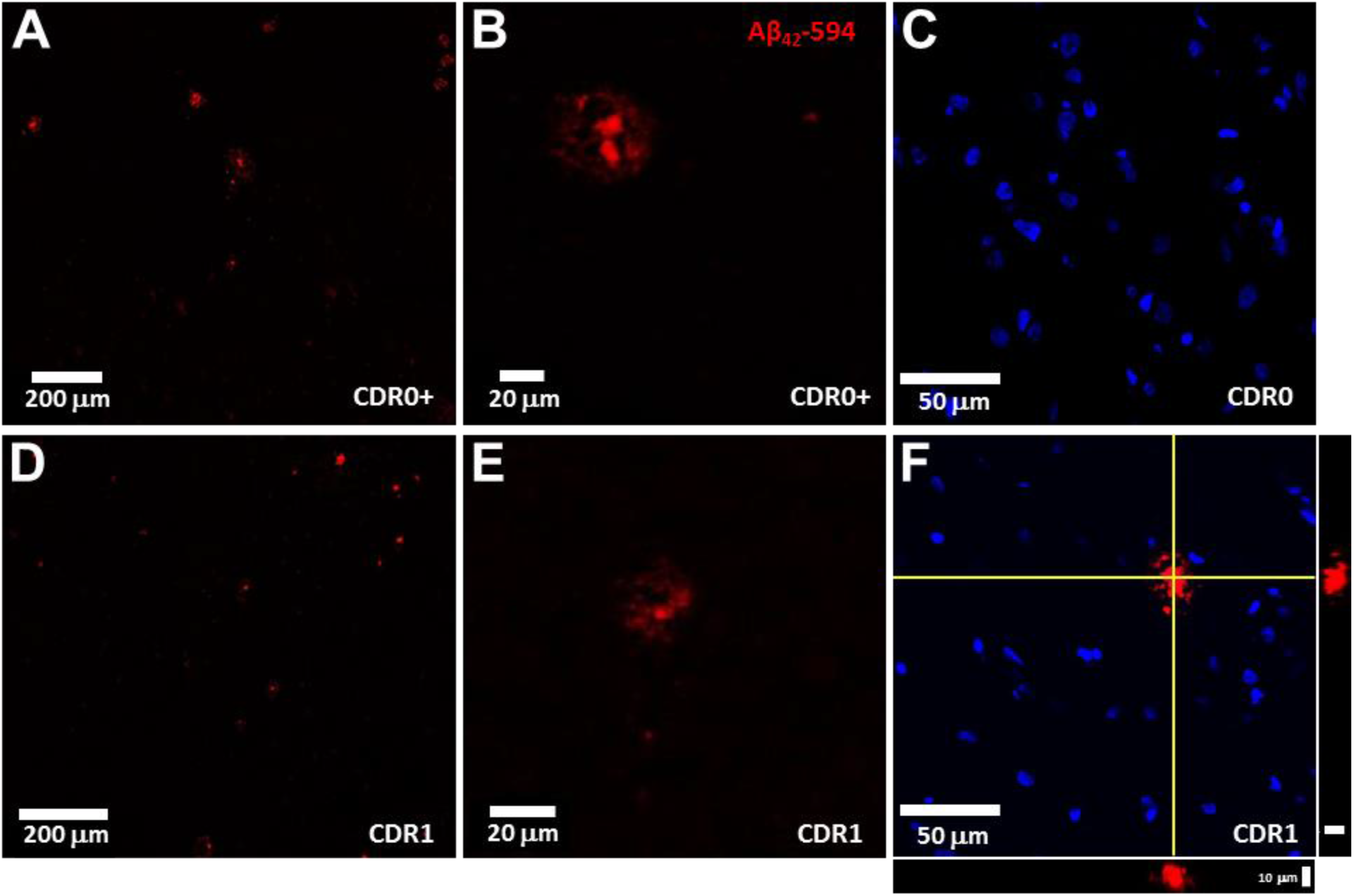
Exemplar fluorescent confocal microscopy of Aβ_1-42_-biotin binding to unfixed, frozen frontal cortex sections. (**A, D**) Fluorescent confocal images of labelled (streptavidin-Alexa594, red channel) Aβ_1-42_-biotin binding in elderly high-pathology controls (CDR0 + plaques) and elderly subjects with mild ADD (CDR1) type representative of those used for quantitation. Scale bar = 200 μm. (**B, E**) Higher magnification confocal images reveal distinct plaque morphology and minimal background fluorescence in both in elderly high-pathology controls (CDR0 + plaques) and elderly subjects with mild dementia of the Alzheimer (CDR1) type. Scale bar = 20 μm. (**C**) Fluorescent images of labelled (streptavidin-Alexa594, red channel) Aβ_1-42_-biotin binding display an absence of plaque structure morphology with minimal background signal in cognitively normal elderly subjects without plaque pathology. Nuclei stained with DAPI (blue channel). Scale bar = 50 μm. (**F**) Fluorescent images of labelled (streptavidin-Alexa594, red channel) Aβ_1-42_-biotin binding display punctate staining of a central core with peripheral decoration in elderly subjects with mild dementia of the Alzheimer (CDR1) type. Nuclei stained with DAPI (blue channel). Scale bar = 50 μm. Orthogonal XZ and YZ views, centered on the yellow crosshairs, demonstrate the labelling extent through the tissue section. Scale bar = 10 μm.

**Figure 3:**
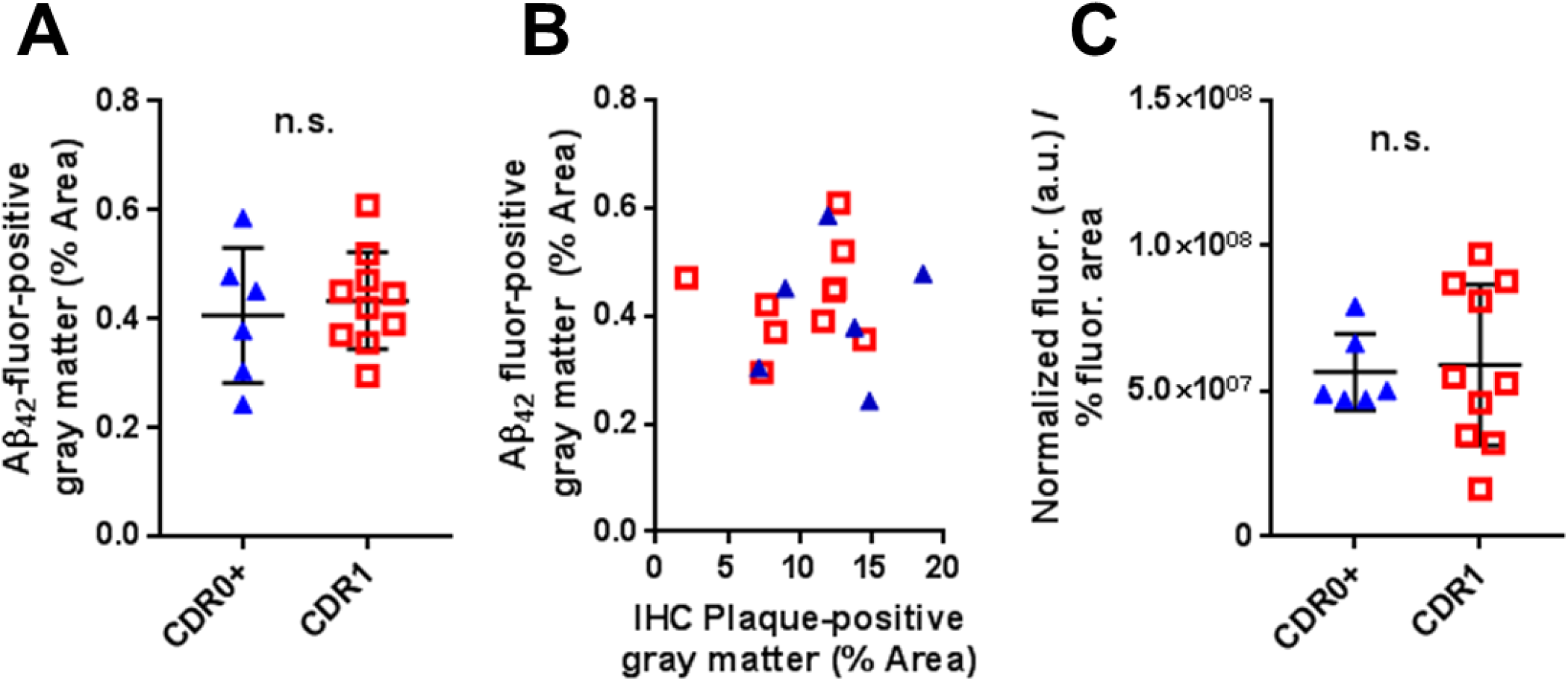
Assessments based on confocal fluorescence microscopy of Aβ_1-42_-biotin binding does not distinguish between elderly high-pathology controls and elderly subjects with mild dementia of the Alzheimer type. (**A**) Gray matter coverage of fluorescently labelled (streptavidin-Alexa594) Aβ_1-42_-biotin binding in CDR 0 + plaques group versus CDR 1 group (not significant [n.s.] by Mann–Whitney U test). (**B**) Correlations between fluorescently labelled Aβ1-42-biotin gray matter coverage and overall Aβ plaque coverage. (**C**) Ratio of the normalized fluorescently labelled Aβ_1-42_-biotin signal to the percentage fluorescent positive coverage (not significant [n.s.] by Mann–Whitney U test).

### No difference between binding of biotinylated Aβ_1-42_ bound to slices from nondemented controls vs. slices from patients with early ADD: results from ELISA

Confocal imaging is most sensitive to tightly localized fluorescence, such as that due to binding of Aβ to discrete plaques and peri-plaque structures. However, it is also possible that other brain tissues could bind and buffer Aβ more diffusely, which might be difficult to detect using confocal microscopy. Therefore, we also used a parallel biochemical approach in which we incubated separate sets of frozen brain slices from the same patients with biotinylated Aβ_1-42_, washed, and then lysed the slices in formic acid to solubilize all of the Aβ in the slices for measurement by ELISA. In agreement with the previous results, we found no difference between the biotinylated Aβ_1-42_ in the lysates from slices from nondemented controls vs. slices from patients with early ADD (**Fig 4A**, p=0.0559). There were no differences between groups after normalizing by total Aβ_1-x_ in the lysates (**Fig 4B**, p=0.7925), nor after normalizing by Aβ plaque coverage (**Fig 4C**, p=0.0934). Thus, an orthogonal approach to measurement of buffering capacity similarly did not differentiate demented from non-demented patients.

**Figure 4:**
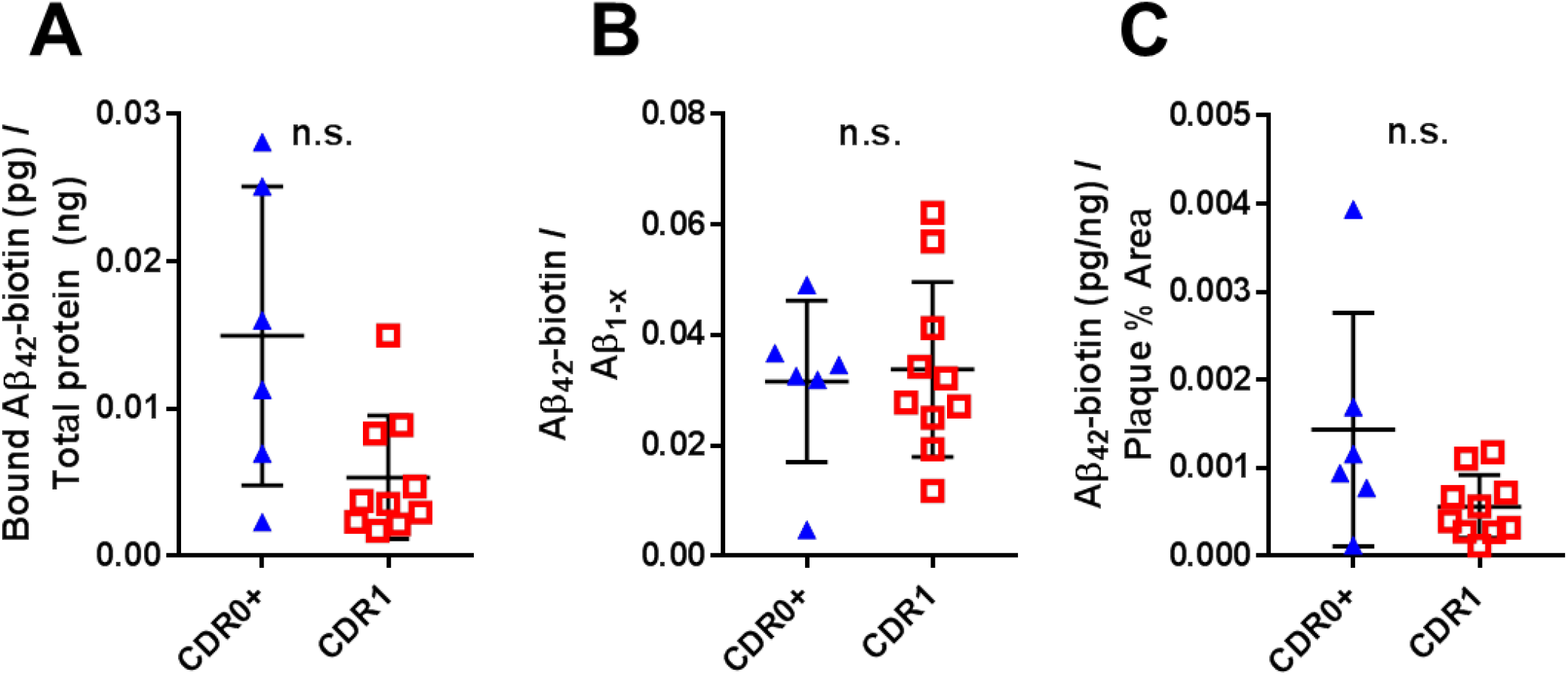
Assessments based on enzyme-linked immunosorbent assay of bound Aβ_1-42_-biotin does not distinguish between elderly high-pathology controls and elderly subjects with mild dementia of the Alzheimer type. (**A**) There was no difference between groups in the levels of overall Aβ_1-42_-biotin recovered from dissociated tissue, as measured using an indirect ELISA which only detects biotinylated Aβ (not significant [n.s.] by Mann–Whitney U test). Data expressed as picograms Aβ per nanogram of total measurable protein. (**B**) The ratio of the amount of Aβ_1-42_-biotin to the amount of Aβ_1-x_ as measured by sandwich ELISA was not different between groups (not significant [n.s.] by Mann–Whitney U test). (**C**) The ratio of Aβ Aβ_1-42_-biotin as measured by sandwich ELISA to the percent gray matter plaque coverage did not differ between groups (not significant [n.s.] by Mann–Whitney U test).

### Synthetic Aβ_1-42_ is partially aggregated at baseline and aggregates further over time during the binding experiments

Synthetic Aβ_1-42_ can be induced to form a large number of aggregation forms, from dimers to higher order oligomers. We performed size exclusion chromatography under neutral pH conditions to examine the size forms of the synthetic Aβ_1-42_ in the solutions that were applied to the slices, and again to examine the size forms present in the supernatant from the slices after the 18 hour incubation on slices at room temperature. At baseline, most of the Aβ_1-42_ appeared monomeric, running in fractions 18 to 22 between the 17kDa and 1.35kDa size standards (**Fig 5** – black solid circles). A small portion appeared to be high molecular weight running in fractions 8 to 9 equivalent to the 670kDa size standards. After incubation, the Aβ was still mainly monomeric, but a portion had shifted to slightly larger appearing forms, in fractions 16 to 18 between the 44kDa and 17kDa size standards (**Fig 5** – red open circles).

**Figure 5:**
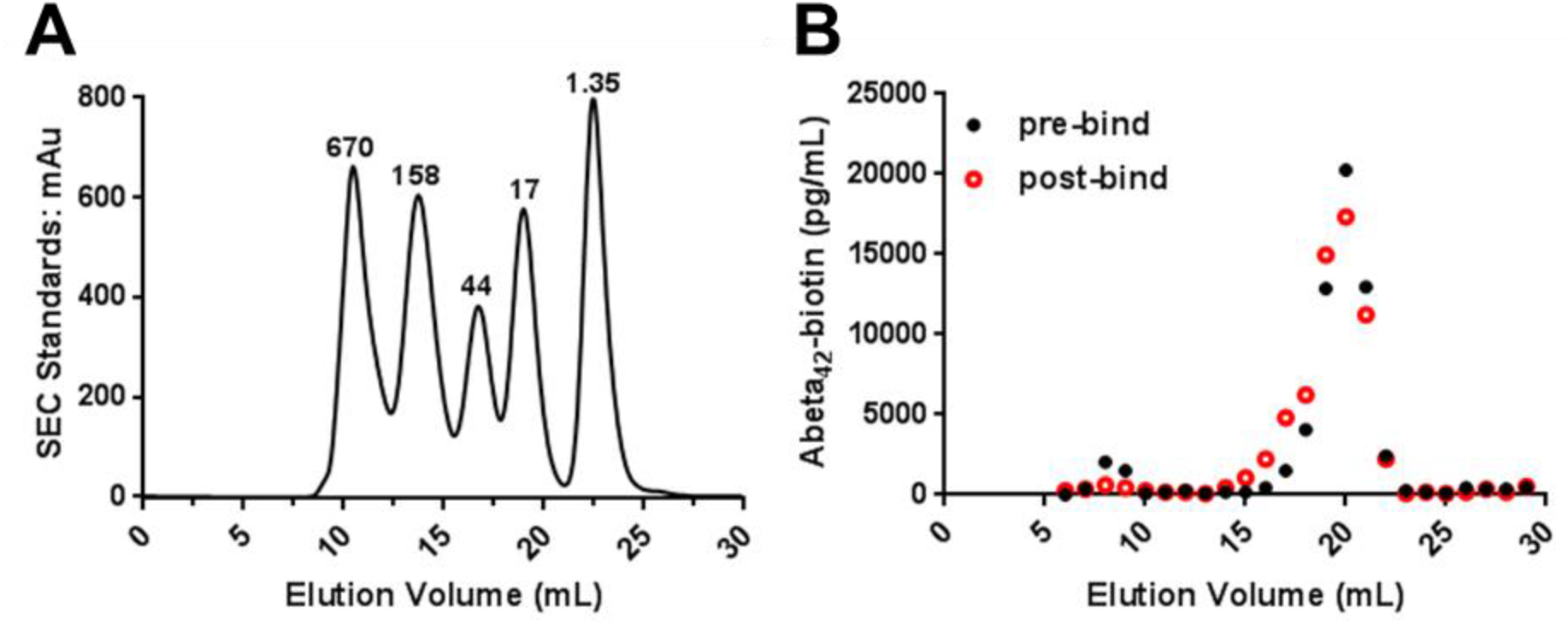
Size exclusion chromatography reveals that the Aβ_1-42_-biotin monomer remained predominantly low molecular weight during the experiment. (**A**) Globular protein molecular weight standards as run on a size exclusion Superdex 200 10/300 GL column. mAu = milli-absorbance units, kDa = kilodaltons. (**B**) Aβ_1-42_-biotin assessed by indirect enzyme-linked immunosorbent assay (ELISA) in the initial incubation condition (black closed circles) and the post overnight incubation condition (red open circles). The predominance of the Aβ_1-42_-biotin elutes as a low molecular weight peak in fractions 18 to 21 likely corresponding to monomer, and a minority of the Aβ_1-42_-biotin eluted as a high molecular weight peak in fractions 8 and 9 corresponding to aggregated Aβ. The post overnight incubation sample has a minor shift to include a shoulder of fractions 16 and 17 in the low molecular weight peak suggesting the accumulation of oligomeric species while most the sample remained monomeric.

## Discussion

In summary, synthetic biotinylated Aβ_1-42_ binds to plaque-like structures indistinguishably in frozen brain slices from demented vs. non-demented high Aβ plaque pathology donors. This negative result does not support the hypothesis that a generalizable differential buffering of Aβ by plaques underlies the difference between demented and non-demented high plaque pathology patients. However, appropriate methods to definitively address the plaque buffering hypothesis have not yet been developed. Most importantly, native soluble Aβ aggregates may have very different properties from synthetic biotinylated Aβ_1-42_ due to differences in proteoforms(9), aggregation states, and potentially associated proteins (2).

### Relationship to Previous Studies

From a methodological perspective, our approach has similarities to those used previously by others. For example, Guo et al.(10) demonstrated that exogenously applied fluorescently labelled Aβ_1-42_ bound plaques in unfixed frozen human cortical sections, though their binding assays were performed for 30 minutes at 37°C rather than over 18 hours at room temperature as in our studies. Esler et al. (11) reported that synthetic Aβ exposed to human cortical sections for a short time unbound quickly upon washing, whereas synthetic Aβ exposed for longer times unbound at substantially slower rate. These results, and others, led to the two process ‘dock-lock’ model of soluble Aβ interacting with plaques. We specifically used an 18 hour binding time to attempt to assess both processes in our studies. Tseng et al.(12) reported that while monomeric Aβ_1-40_ bound plaques from AD brain, aged high molecular weight synthetic Aβ aggregates did not show detectible plaque binding. Concordantly, the majority of the Aβ in our experiments was monomeric throughout the experiment, though there appeared to be some aggregation over time. However, none of these previous investigations compared Aβ binding to slices from brains of patients with dementia vs. from patients with indistinguishable plaque burden but no dementia.

The above discussed studies involved binding to *ex vivo* human brain tissue plaques, whereas other groups have investigated similar properties in living animal models. For example, Gureviciene et al. (13) infused fluorescently labelled monomeric Aβ into APPswe/PSEN1dE9 mice and measured incorporation of the Aβ by two-photon excitation fluorescence microscopy. They reported accumulation of fluorescent Aβ preferentially around plaques compared to other brain regions. This result was interpreted as consistent with plaque buffering of soluble Aβ *in vivo*.

### Limitations and Future Directions

The main limitation of our approach was that we used synthetic monomeric Aβ. Future studies will be required to assess the properties of human brain-derived Aβ monomers and soluble Aβ aggregates. Importantly, the characteristics of native human brain-derived Aβ may not be comparable to those of synthetic Aβ aggregates due to the wide variety of proteoforms, aggregation states, and potentially co-associated proteins. For example, Jin et al.(14) demonstrated that synthetic Aβ dimers are orders of magnitude less biologically active than human brain-derived Aβ immunoreactive species of comparable apparent size. Similarly, Noguchi et al (15) reported that native human brain high molecular mass Aβ aggregates were substantially more toxic than similar size synthetic Aβ aggregates.

The ability to label Aβ aggregates extracted and purified from the human brain without disrupting their native tertiary structure will be essential for these future endeavors. Having methods to compare the initial state and the labelled state will be important to ensure that experiments can be interpreted accurately. We envision testing for whether the process of labeling native brain Aβ aggregates disrupts their properties by using a competition assay; for example if the directly measured affinity of labeled native brain Aβ is similar to the inferred affinity of unlabeled native brain Aβ as a competitor, this would be reassuring.

Ongoing work in our group and that of others should soon begin to address hypothesis 2 above by directly determining the Aβ proteoform composition of soluble Aβ aggregates extracted from the brains of patients with dementia vs. patients with indistinguishable plaque burden but no dementia. From there, the ability of plaques to buffer soluble Aβ aggregates from the two patient sources or synthetic versions of differentially abundant proteoforms will be tested directly.

A second limitation involves the use of frozen *ex vivo* brain slices. If active cellular processes play a role in the functional buffering of Aβ, this would not be detectible in our assays. In theory, microdose PET tracer labeled Aβ could be infused into local regions in the brains of patients undergoing other clinically indicated neurosurgical procedures such as shunting for normal pressure hydrocephalus. Retention vs. clearance of the PET tracer label could be then compared between non-demented low plaque, non-demented high plaque, and demented high plaque individuals.

A third limitation is that we did not measure the full kinetic binding properties of the Aβ in our experiments, nor did we measure binding at physiological temperature. It is possible that differences between the two groups in the kinetics of Aβ binding could be important, even though the likely near-equilibrium binding we measured did not differ between groups.

### Implications

A more complete understanding of the differences between the Aβ found in the brains of patients with dementia vs. patients with equivalent presence of plaque burden but no dementia may be key to developing Aβ-targeted therapeutics for ADD. It is quite possible that despite many years and many clinical trials, the appropriate Aβ therapeutic targets most relevant to dementia still have not been identified.

## Acknowledgements

We would like to thank the participants and their families of the Knight ADRC who donated their brain tissue for medical research. We would also like to thank Dennis Oakley for technical assistance with confocal microscopy, Emeka Fountain for technical assistance, David Holtzman and Hong Jiang for antibodies, the Hope Center Alafi Neuroimaging Lab for access to the NanoZoomer microscope, and the Washington University Center for Cellular Imaging for access to the confocal microscope. We gratefully acknowledge helpful discussions with John Maggio, Tom Ellenberger, Paul Kotzbauer, and John Cirrito.

## Author Contributions

T.J.E. and D.L.B designed experiments. T.J.E. performed experiments. T.J.E. and D.L.B. performed primary data analysis. M.G. contributed novel analytical methods. N.J.C. provided brain tissue and pathological analyses. D.L.B. obtained funding. T.J.E and D.L.B. wrote the manuscript with intellectual contributions from all authors.

## References

1. Day GS, Gordon BA, Jackson K, Christensen JJ, Rosana Ponisio M, Su Y, et al. Tau-PET Binding Distinguishes Patients With Early-stage Posterior Cortical Atrophy From Amnestic Alzheimer Disease Dementia. Alzheimer Dis Assoc Disord. 2017;31(2):87–93.

2. Brody DL, Jiang H, Wildburger N, Esparza TJ. Non-canonical soluble amyloid-beta aggregates and plaque buffering: controversies and future directions for target discovery in Alzheimer's disease. Alzheimer's research&therapy. 2017;9(1):62.

3. Esparza TJ, Zhao H, Cirrito JR, Cairns NJ, Bateman RJ, Holtzman DM, et al. Amyloid-beta oligomerization in Alzheimer dementia versus high-pathology controls. Ann Neurol. 2013;73(1):104–19.

4. Koffie RM, Meyer-Luehmann M, Hashimoto T, Adams KW, Mielke ML, Garcia-Alloza M, et al. Oligomeric amyloid beta associates with postsynaptic densities and correlates with excitatory synapse loss near senile plaques. Proc Natl Acad Sci U S A. 2009;106(10):4012–7.

5. Koffie RM, Hashimoto T, Tai HC, Kay KR, Serrano-Pozo A, Joyner D, et al. Apolipoprotein E4 effects in Alzheimer's disease are mediated by synaptotoxic oligomeric amyloid-beta. Brain: a journal of neurology. 2012;135(Pt 7):2155–68.

6. DeKosky ST, Scheff SW. Synapse loss in frontal cortex biopsies in Alzheimer's disease: correlation with cognitive severity. Ann Neurol. 1990;27(5):457–64.

7. Terry RD, Masliah E, Salmon DP, Butters N, DeTeresa R, Hill R, et al. Physical basis of cognitive alterations in Alzheimer's disease: synapse loss is the major correlate of cognitive impairment. Ann Neurol. 1991;30(4):572–80.

8. Esparza TJ, Wildburger NC, Jiang H, Gangolli M, Cairns NJ, Bateman RJ, et al. Soluble Amyloid-beta Aggregates from Human Alzheimer's Disease Brains. Sci Rep. 2016;6:38187.

9. Wildburger NC, Esparza TJ, LeDuc RD, Fellers RT, Thomas PM, Cairns NJ, et al. Diversity of Amyloid-beta Proteoforms in the Alzheimer's Disease Brain. Scientific reports. 2017;7(1):9520.

10. Guo JP, Yu S, McGeer PL. Simple in vitro assays to identify amyloid-beta aggregation blockers for Alzheimer's disease therapy. J Alzheimers Dis. 2010;19(4):1359–70.

11. Esler WP, Stimson ER, Jennings JM, Vinters HV, Ghilardi JR, Lee JP, et al. Alzheimer's disease amyloid propagation by a template-dependent dock-lock mechanism. Biochemistry. 2000;39(21):6288–95.

12. Tseng BP, Esler WP, Clish CB, Stimson ER, Ghilardi JR, Vinters HV, et al. Deposition of monomeric, not oligomeric, Abeta mediates growth of Alzheimer's disease amyloid plaques in human brain preparations. Biochemistry. 1999;38(32):10424–31.

13. Gureviciene I, Gurevicius K, Mugantseva E, Kislin M, Khiroug L, Tanila H. Amyloid Plaques Show Binding Capacity of Exogenous Injected Amyloid-beta. J Alzheimers Dis. 2017;55(1):147–57.

14. Jin M, Shepardson N, Yang T, Chen G, Walsh D, Selkoe DJ. Soluble amyloid beta-protein dimers isolated from Alzheimer cortex directly induce Tau hyperphosphorylation and neuritic degeneration. Proc Natl Acad Sci U S A. 2011;108(14):5819–24.

15. Noguchi A, Matsumura S, Dezawa M, Tada M, Yanazawa M, Ito A, et al. Isolation and characterization of patient-derived, toxic, high mass amyloid beta-protein (Abeta) assembly from Alzheimer disease brains. J Biol Chem. 2009;284(47):32895–905.

